# The development of Bayesian integration in sensorimotor estimation

**DOI:** 10.1101/136267

**Authors:** Claire Chambers, Taegh Sokhey, Deborah Gaebler-Spira, Konrad Kording

## Abstract

If the brain is inherently Bayesian, then behavior should show the signatures of Bayesian computation from an early stage in life without the need for learning. Children should integrate probabilistic information from prior and likelihood distributions to reach decisions and should be as statistically efficient as adults. To test this idea, we examined the integration of prior and likelihood information in a simple position estimation task comparing children aged 6-11 years and adults. During development, estimation performance became closer to the statistical optimum. Children use likelihood information as well as adults but are limited in their use of priors. This finding suggests that Bayesian behavior is not inherent but learnt over the course of development.

---

The behavior of adults under uncertainty is well described by Bayesian inference in sensorimotor behavior (Kording and Wolpert, 2004), perception (Knill and Richards, 1996), cognition and reasoning tasks (Tenenbaum et al., 2011), and cue combination across and within sensory modalities (Ernst and Banks, 2002). Adult humans seem to integrate information in a way predicted by Bayesian statistics.

Findings of Bayesian behavior have led to the theory that the underlying neural computations are Bayesian. It has been argued that neural activity reflects probabilistic population codes that directly implement Bayesian computations (Ma et al., 2006). However, Bayesian behavior simply represents optimal behavior under uncertainty and there are ways of generating optimal behavior that do not explicitly implement Bayesian computation (Rao, 2004). Therefore, previous research has not fully established whether the neural code is inherently Bayesian.

If neural circuits are evolved to implement Bayesian computations, then behavior should always show Bayesian signatures. Therefore, children too should act in accordance with the rules of Bayesian integration. Specifically, they should weigh information according to its relative uncertainty in simple tasks.

Indeed, a good number papers ask how Bayesian children are. Looking times in infants are consistent with early optimal integration (Téglás et al., 2011) and young children use probabilistic information to infer causality (Kushnir and Gopnik, 2007). However, in the former case measurements are indirect and in the latter case predictions can only be qualitative. However, work on cue combination and integration of value-based information shows that children do not integrate as adults do until 9-11 years (Dekker et al., 2015; Dekker and Nardini, 2016; Gori et al., 2008; Nardini et al., 2010, 2008). Therefore, it is unclear whether the behavior of children is consistent with use of Bayesian inference.

Here we investigate whether integration of current with past information to perform actions under uncertainty is present in young children or is acquired over the course of development. In our paradigm, we examine the use of probabilistic information to perform a simple sensorimotor estimation task (Berniker et al., 2010; Kording and Wolpert, 2004; Vilares et al., 2012). Visual targets were drawn from a prior distribution and participants were shown uncertain sensory information. While all age groups learned to use the uncertainty of sensory information, children did not exploit the uncertainty of the prior to perform estimations as adults did, with this gradually emerging during development.

## Methods

### Experimental details

We aimed to examine probabilistic inference during sensorimotor estimation in a child population. Our task was designed to examine use of probabilistic information during sensorimotor estimation (Fig. 1a, Acuna, Berniker, Fernandes, & Kording, 2015; Berniker et al., 2010; Vilares et al., 2012). Previous findings indicate that adults weigh information according to its reliability and learn priors in a manner which resembles Bayesian integration during sensorimotor estimation. For the purposes of the current study, we adapted the experimental protocol for child participants, by using a concept that was engaging to children, using simplified instructions, and by reducing the number of trials.

**Figure 1.**
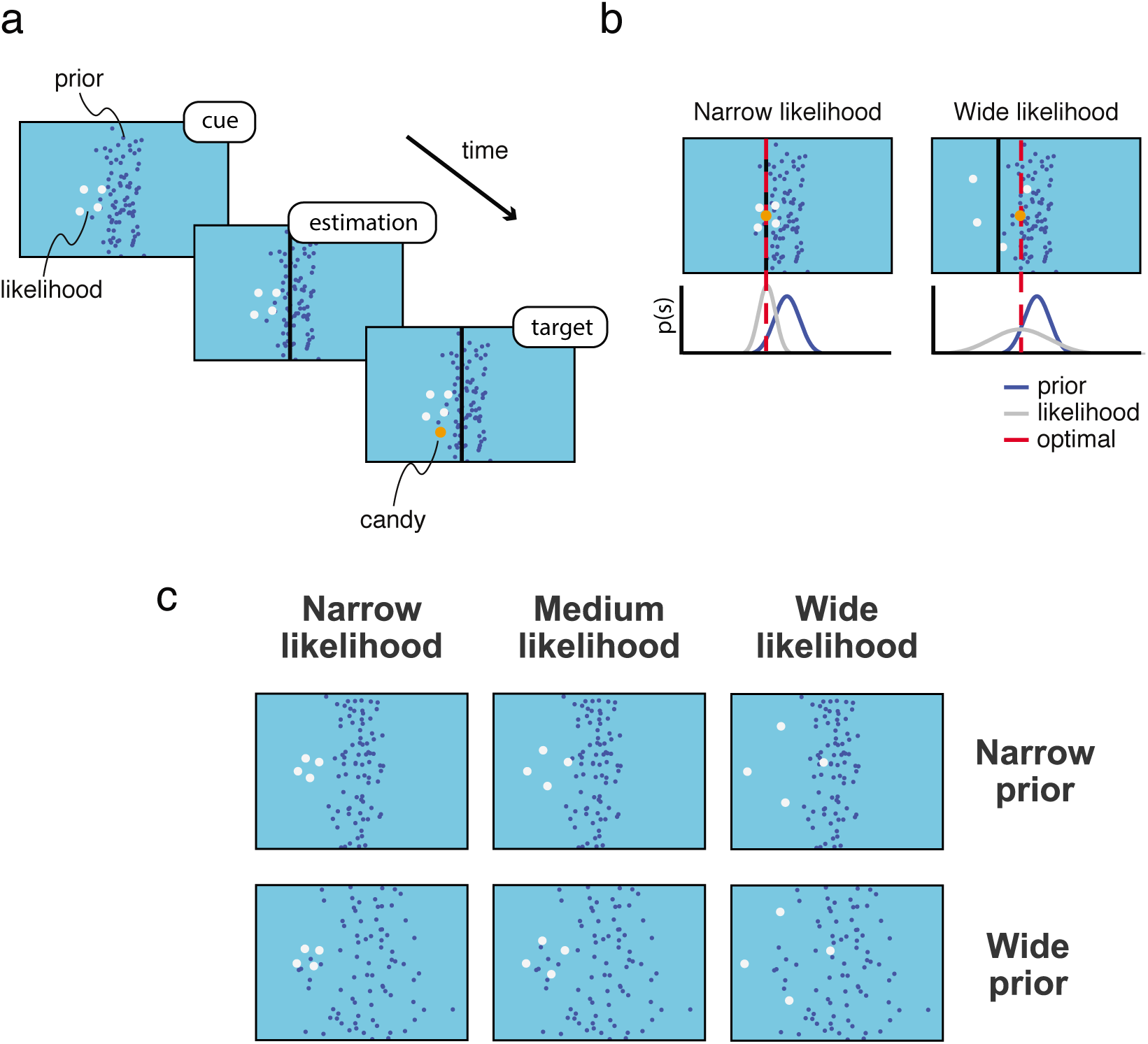
(**a**) Experimental protocol. Participants were shown a visual cue (likelihood) with experimentally controlled uncertainty (splash), created by a hidden target (candy) drawn from a prior distribution. Participants were told that the splash was created by candy falling into a pond. Participants were prompted to place a vertical bar (net) where the hidden target fell, and were then shown feedback on target location. (**b**) Relying on the likelihood. A simple strategy would be to rely entirely on likelihood information by pointing at its centroid on each trial. While this strategy is close to optimal when the likelihood is precise or narrow (black bar, left panel), this strategy is less successful when the likelihood is wider (black bar, right panel), as samples from the likelihood become a less reliable indicator of target location and the optimal estimate shifts closer to the prior mean. The optimal strategy involves weighing prior and likelihood information according to their relative uncertainties (**c**) Experimental design. In order to quantify integration of the prior and likelihood, we measured reliance on the likelihood (*Estimation slope*) under different conditions of prior variance and likelihood variance. The prior could be narrow or wide, and the likelihood could be narrow, medium, or wide.

Participants were 16 children (8 males) aged 6-8 years (M=6.94, SD=0.77), 17 children (8 males) aged 9-11 years (M=10.06, SD=0.75), and 11 adults (5 males) aged over 18 years (M=27.27, SD=5.31). The data of four participants aged 5 years were excluded due to difficulty in using a computer mouse. The data of two additional participants were excluded due to looking away from the screen during the experiment.

In a quiet room, participants sat in front of a 52 cm wide, 32.5 cm high computer monitor. Before starting the experiment, we presented participants with the instructions that someone behind them was throwing candy into a pond, represented by the screen; and that their aim was to estimate where the candy target landed and catch as many candy as possible over the course of the experiment. Candy targets were drawn from a Gaussian distribution centered at the middle of the screen, 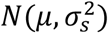. On each trial, they were presented with an uncertain “splash” stimulus for one second and were told that the splash was caused by a hidden candy target. The splash was *n*=4 samples from a Gaussian likelihood distribution that was centered on target location 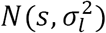. Participants provided an estimate of the candy target’s location on the horizontal axis using a vertical bar that extended from the top to the bottom of the screen. The net appeared at the same time as the splash at a random location on screen. Participants had 6 seconds to respond. After providing a response, they were shown the true candy location.

One simple strategy for performing sensorimotor estimation under uncertainty is to consistently judge target location at the center of the splash – i.e. full reliance on the likelihood. This strategy works well when the likelihood distribution is narrow, because the closely-spaced points of the splash are an accurate indicator of target location (Fig. 1b, left). However, full reliance on the likelihood would cause a participant to miss targets more frequently as the likelihood distribution widens (Fig. 1b, right). When sensory information is unreliable, rather than relying on the likelihood completely, we maximize performance by giving more weight to our prior belief on target location. More generally, the best or optimal strategy involves weighing sources of information according to their relative precision.

Formally, weighing sources of information according to their relative precision corresponds to Bayesian inference. An optimal Bayesian observer combines noisy sensory information from the likelihood, 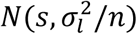 with their learned prior, 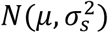, resulting in a posterior distribution over target location, 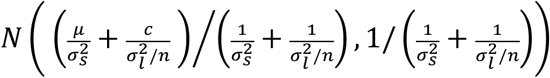. The mean of the posterior is a mean of the prior and centroid of the likelihood, *c*, weighted by their precisions. From this posterior distribution, an estimation, 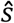, is computed. Therefore, the optimal reliance on the likelihood is a function of prior and likelihood uncertainties, 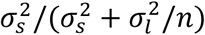. We can manipulate the prior and likelihood variance and measure their influence on participants’ reliance on the likelihood, in order to investigate probabilistic information during sensorimotor estimation.

To investigate how children use probabilistic information during sensorimotor estimation, we manipulated the variances of prior distribution and likelihood distributions. We used a Gaussian prior distribution with a mean at the center of the screen and standard deviation of .03 (Narrow Prior) or .1 (Wide Prior) in units of screen width. The likelihood distribution was centered on target location and could have a standard deviation of .05 (Narrow Likelihood), .1 (Medium Likelihood), or .25 (Wide Likelihood) in units of screen width. There were six conditions: Narrow Prior – Narrow Likelihood, Narrow Prior – Medium Likelihood, Narrow Prior – Wide Likelihood, Wide Prior – Narrow Likelihood, Wide Prior – Medium Likelihood, and Wide Prior – Wide Likelihood.

The experiment consisted of four blocks, each lasting 120 trials, preceded by a practice block lasting 10 trials. Trials were blocked by prior condition, with all likelihood conditions being presented in randomized order within one block. The prior over target location switched from block to block with a randomly chosen starting condition for each participant (i.e., narrow-wide-narrow-wide or wide-narrow-wide-narrow).

We introduced a number of modifications to engage child participants in the task. Participants were shown how much candy they had won on screen and participants won a “bonus” piece of candy for every ten candy they caught. Sounds were presented to signal successfully catching a target and missed responses when they did not respond within the 6-second time window. Step-by-step instructions were shown to participants on screen before the experiment, to ensure that all participants received the same instructions. Participants were told that their payment was based on the number of candy that they caught.

Ethical approval was provided by the NU IRB #20142500001072 (Northwestern University, USA). Participants signed a consent form before participation. For participants aged under 18 years, a parent provided consent for their child to take part and completed the Developmental Coordination Disorder questionnaire (Wilson et al., 2009), a modified Vanderbilt questionnaire to assess for ADHD (Wolraich et al., 2003), and the Behavior Assessment System for Children, BASC-3, parent rating scales (Reynolds, 2004). After the participant had completed the game, we administered the child mini-mental state evaluation to obtain an approximate assessment of cognitive ability (Ouvrier et al., 1993). No data was excluded on the basis of neuropsychological test results.

### Data analysis

We were interested in the integration of probabilistic information from a prior distribution and sensory information from the likelihood in sensorimotor estimations. To investigate this, we examined whether samples from the likelihood distribution,

*X* = {*x*_1,_*x*_2,_*x*_3_,*x*_4_}, were combined with information about the prior distribution, 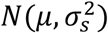, when producing an estimate of target location, 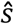. We quantified this for each condition using the extent to which participants relied on the likelihood, given by the linear relationship between the centroid of the splash, 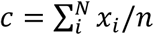, and their estimate on each trial, 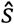. We performed a linear regression with estimations, 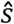, as dependent variable and the likelihood centroid, *c*, as the independent variable, which resulted in a measure of reliance on the likelihood, which we term the *Estimation slope* (*ES*). We assumed that participants accurately learned the mean of the prior, and set the intercept to 0. If participants relied only on the likelihood to generate their estimate, then they should point close to the centroid of the splash, *c*, on all trials, leading to an *Estimation slope* ≈ 1. If instead participants ignore the likelihood and instead only use their learnt prior, then their estimates should not depend on the *c*, leading to an *Estimation slope* ≈ 0. Therefore, from participants’ estimations we obtain a measure of their reliance on the likelihood or prior.

The theoretical variance of the likelihood and prior used in the experiment provide optimal values for the *Estimation slope*, i.e. how much participants should rely on the likelihood. For an optimal Bayesian observer, sources of information are weighed according to their relative reliabilities, 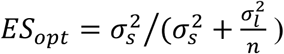. Therefore, in order to quantify how optimal participants were, we can compare *Estimation slope* quantified from participant's data with the optimal *Estimation slope*, *ES*_*opt*_, by computing the absolute difference, |*ES – ES*_*opt*_| in each condition, and then averaging this across conditions, which provided a *Distance to the optimal* score for each participant.

We wanted to know how sensitive participants were to the prior and likelihood variance. We therefore devised separate measures to quantify how much participants distinguished between prior conditions and between likelihood conditions. The sensitivity to the prior was simply the difference between the *Estimation slopes* across prior conditions, which was then summed across likelihood conditions:

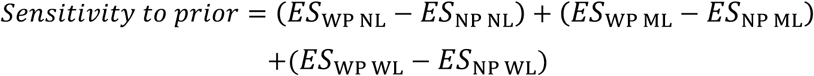

Similarly, the sensitivity to the likelihood was the difference between *Estimation slopes* across likelihood conditions for a fixed prior condition, summed across prior conditions:

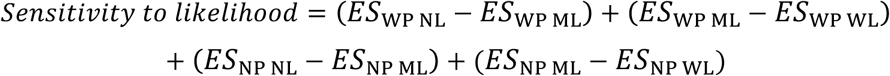

Since these two measures are the signed difference between conditions, a positive score indicates use of statistical information in a way that is predicted by the experimental parameters, whereas a negative score indicates use of statistical information which is opposite to that predicted by the experimental parameters. Together, these two measures account for use of probabilistic information during the task.

To examine subject-specific biases, in the form of an overall tendency to use the likelihood only or prior only regardless of the experimental condition, we computed a *Bias* score for each participant, which was simply the *Estimation slope* averaged across all conditions. This allowed us to examine use of simple response strategies.

To quantify the uncertainty of our estimation of *Estimation slope*, *Distance to optimal*, *Sensitivity to prior*, *Sensitivity to likelihood*, and *Bias*, we performed bootstrapped estimation by resampling the data with replacement 1000 times and performing the fit for each resampled data set. Since the data did not meet the requirements for parametric statistical tests, we performed non-parametric tests on the data: Kruskal-Wallis tests to examine main effects, Mann-Whitney U tests to examine differences between groups, and Wilcoxon signed-rank tests for comparison of individual samples with chance level, Bonferroni-corrected for the number of comparisons.

## Results

We wanted to investigate the development of Bayesian integration. To do so, we examined whether children aged 6-11 years old and adults could learn to use uncertainty of different sources of information (prior and likelihood) during sensorimotor estimation (Fig. 1). We examined use of probabilistic information during development by quantifying participants’ task performance, reliance on the likelihood, the degree to which they deviated from the statistical optimum and their sensitivity to the prior variance and likelihood variance condition.

We first established that all age groups understood and carried out the task, with performance above chance level of 2%. *p(correct)* exceeded chance level for all groups (Table 1). Therefore, differences between age groups cannot be attributed to a lack of understanding of the task.

**Table 1:**
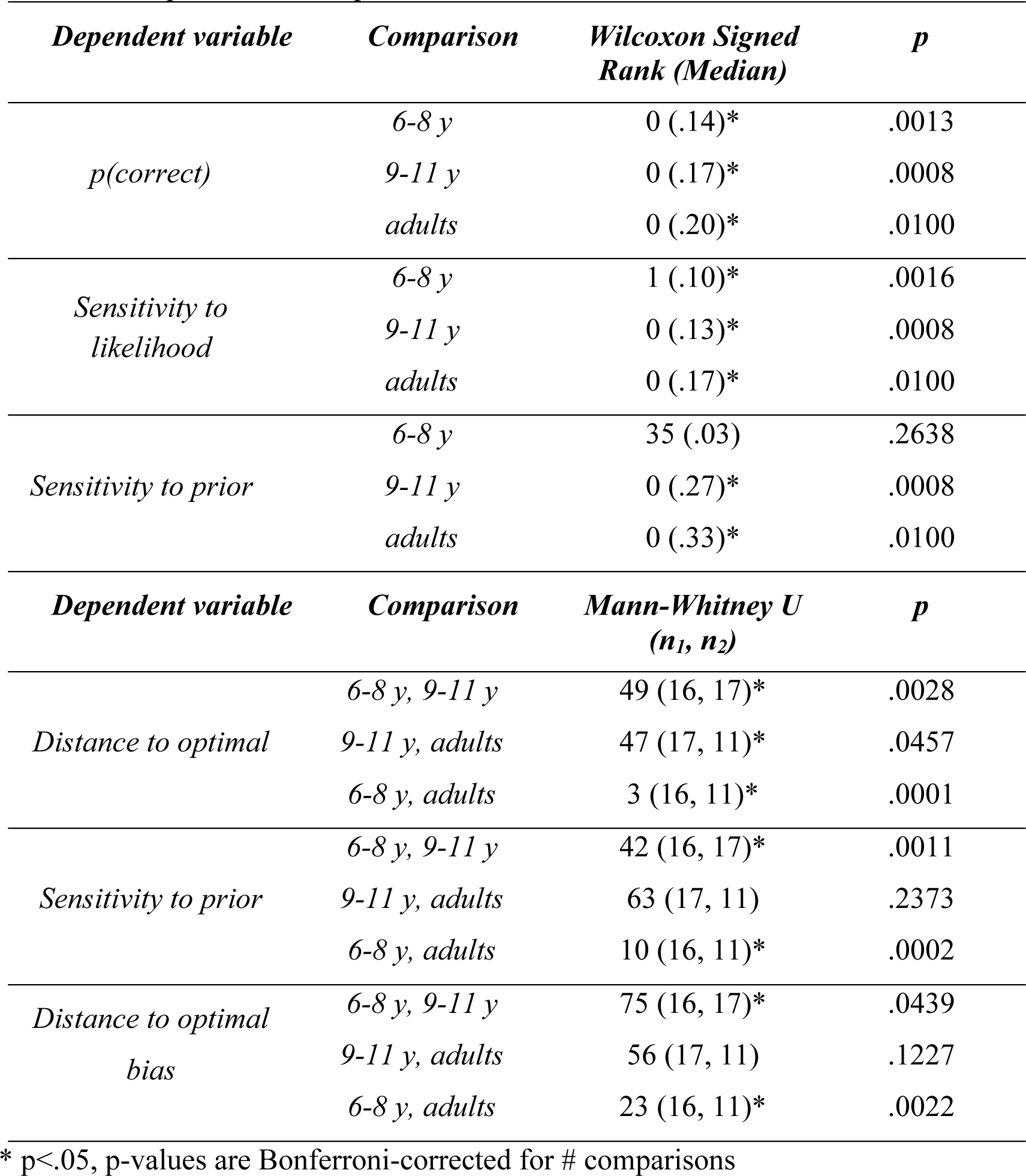
Non-parametric comparisons

In order to investigate sensorimotor estimation under uncertainty, we manipulated the variance of the prior and likelihood and measured the *Estimation slope* in each condition. Our fitting procedure provides reasonable fits to the data for individuals (shown for an 11-year old, Fig. 2a), and across the entire data set (Fig. 2b). This allows us to quantify the nature of prior/likelihood integration for each participant.

**Figure 2.**
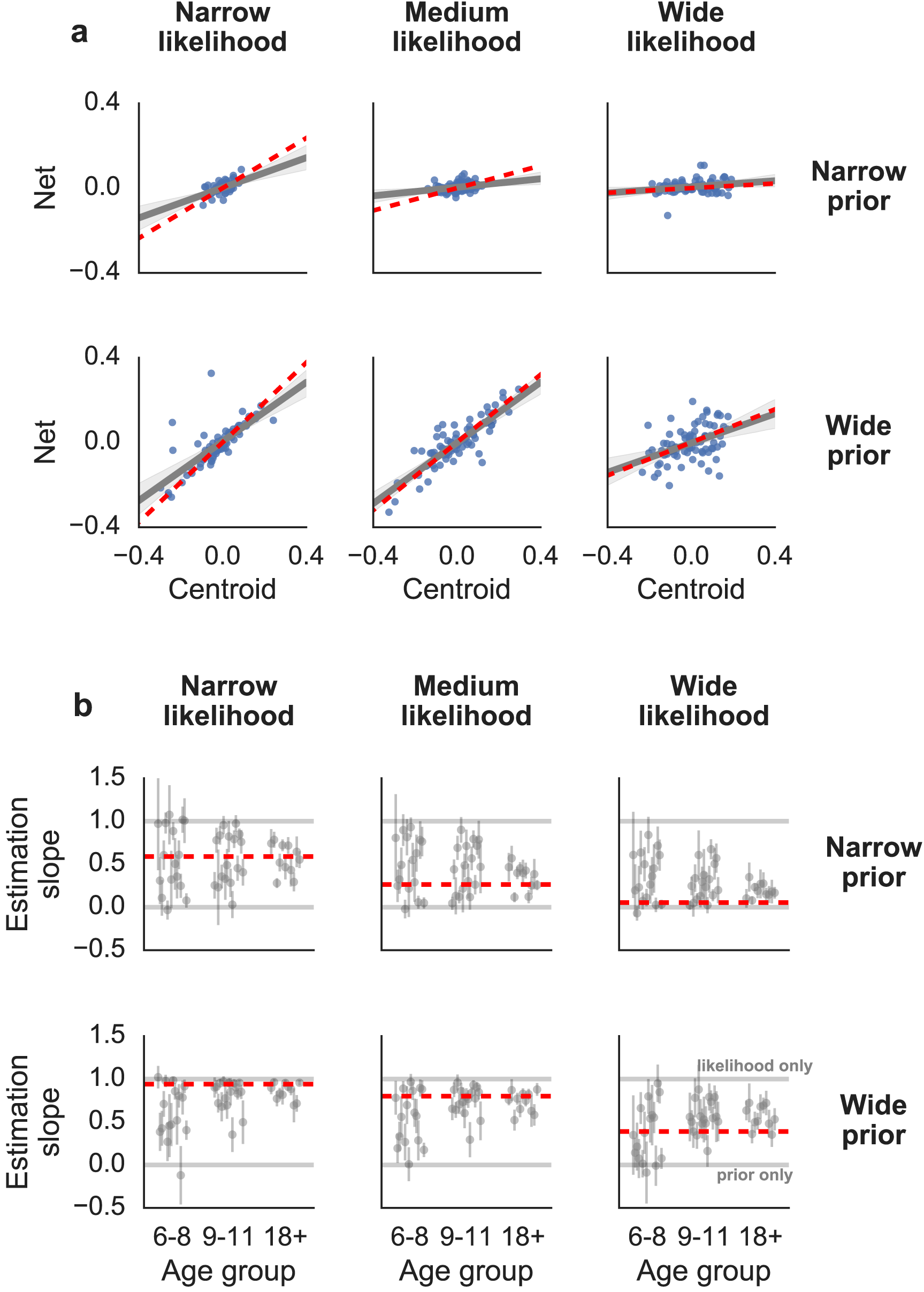
Estimation data. (**a**) Estimation data overlaid with linear fit for a representative participant aged 11 years old. The net position as a function of the centroid of the likelihood is shown for each trial (points). The fitted *Estimation slope* (gray) and the optimal *Estimation slope* (red) are displayed. Each panel displays estimation data for one condition, as defined by prior and likelihood width. (**b**) The median bootstrapped *Estimation slope* of individual participants is shown as a function of age group (error bars = 95% confidence intervals). The optimal *Estimation slope* values are shown (red).

We were interested in whether sensorimotor integration of prior and likelihood improves during development. We, therefore, examined the relationship between the *Estimation slope* measured from the data and the optimal *Estimation slope*, *ES*_opt_. There is a shift toward the statistical optimum during development in each condition (Fig. 2b). For a formal analysis, we computed a distance-to-optimal score for wach participant (Fig. 3b). There was a significant effect of age on *Distance to optimal*, (H(2) = 21.32, p < .0001), with significant differences between all age groups (Table 1). Therefore, sensorimotor estimation shifts toward an optimal integration of prior and likelihood during development.

**Figure 3.**
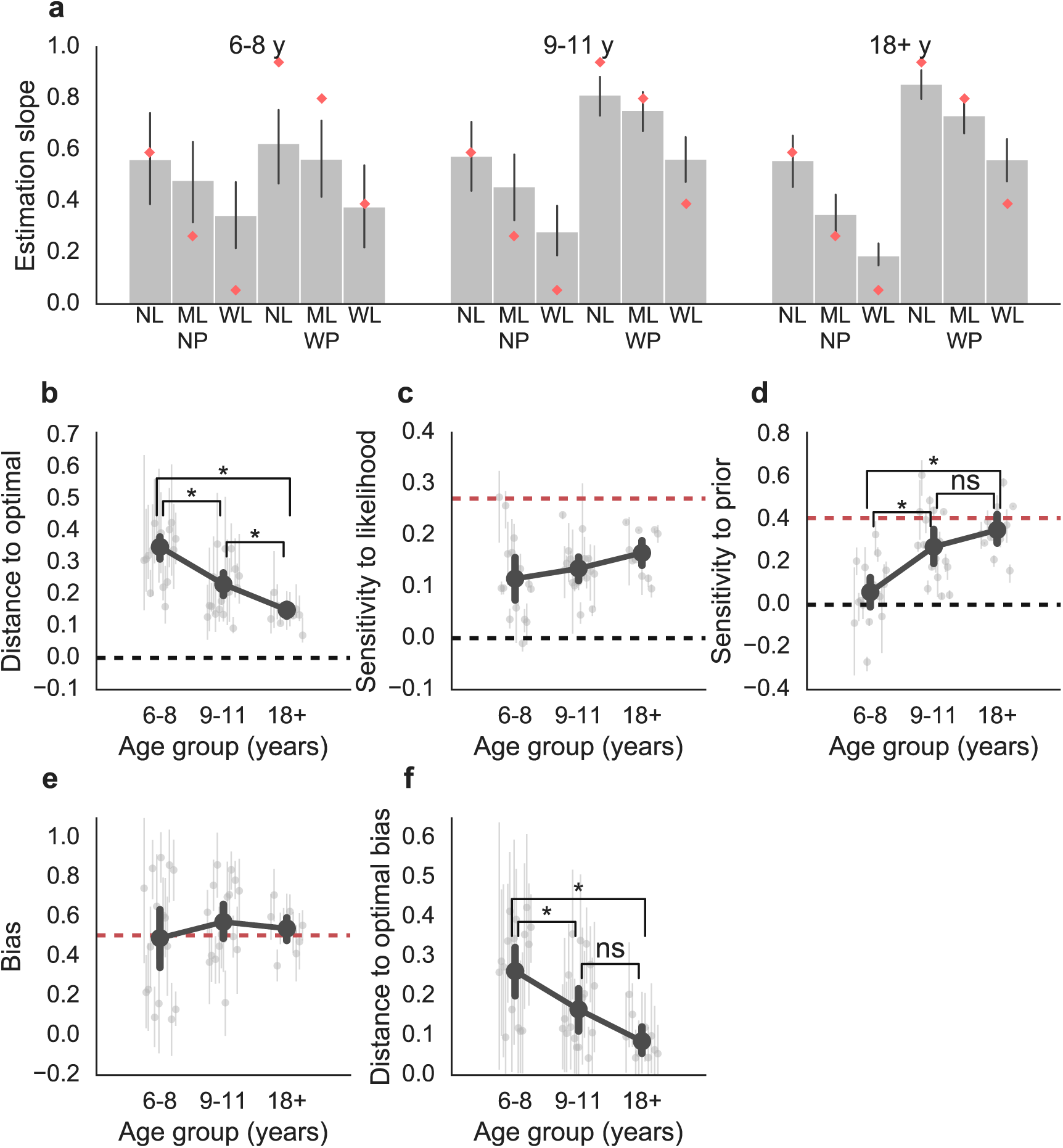
Estimation slope and sensitivity to statistical information and bias. (**a**) Mean *Estimation slope* as a function of prior (NP, WP), likelihood (NL, ML, WL), and age group (error bars = 95% CI). Optimal values are shown (red). (**b**) Distance to the optimal Estimation slope as a function of age group. Here and elsewhere, scores of individual participants are shown in gray overlaid with the mean for each age group (error bars = 95% CI), with optimal values shown in red. *Distance to optimal* decreases with age. (**c**) *Sensitivity to likelihood* was computed as a difference in *Estimation slope* between likelihood conditions. There is no significant effect of age group. (**d**) *Sensitivity to prior* was computed as the difference in *Estimation slope* between prior conditions. This increases with age. (**e**) *Bias* or overall tendency to rely on the likelihood or prior. Younger participants are more biased, but without an overall bias at the group level. (**f**) *Distance to optimal bias*, the absolute difference between a participant’s bias and the optimal bias. This decreases with age.

Two possible contributors to the shift toward the optimal are participants’ use of prior information and their use of likelihood information. We found no evidence for an effect of age on *Sensitivity to likelihood* (Fig. 3c, H(2) = 5.21, p =.0739). *Sensitivity to likelihood* was significantly above zero in all age groups (Table 1). Children aged 6-8 years had already learned to distinguish between likelihood conditions. There was a significant effect of age on *Sensitivity to prior* (Fig. 3d, H(2) = 19.13, p <.0001), with significant differences between age groups, except between 9-11 year olds and adults (Table 1). *Sensitivity to prior* did not exceed chance in children aged 6-8 years, and was above chance in children aged 9-11 years and adults (Table 1). Therefore, even young children can use probabilistic information from the likelihood, but are limited in their use of priors, with this emerging over the course of development.

Young children’s responses may be partly driven by simpler strategies. We therefore quantified the overall bias in participants’ responses. Biases appear to be more prevalent in the responding of individual participants at a younger age, but without an overall bias across participants (Fig. 3e). There is a significant effect of age on *Distance to optimal bias* (H(2) = 11.99, p =.0024, Fig. 3f). Comparisons between age groups were significant except between 9-11 year olds and adults (Table 1). Therefore, simple biases are more prominent in the responding of the youngest participant group.

## Discussion

We examined the development of Bayesian integration during sensorimotor estimation. We found that children consistently deviate from optimal use of statistical information relative to adults. Children used sensory information (likelihood), as adults did. However, the youngest age group (6-8 years) did not distinguish between prior conditions, with use of the prior increasing with age. Young children’s estimations were driven by simple biases, which decreased during development. While young children used the uncertainty of an immediate source of information, they did not incorporate the distribution of targets over several trials into their estimations.

We found that young children aged 6-8 years used the prior to a lesser degree than older children and adults. However it is possible that under different experimental conditions young children may have used the prior, in a longer experiment they may have eventually learned the prior, or if the task required them to actively engage with the prior. Nevertheless, our findings still show that older children and adults use priors in their estimations more readily than children do.

Cognitive explanations can be offered as to why young children fail to incorporate the prior in their estimations. Acquiring the prior involves integrating information over a longer timescale. Therefore, possible contributors to increased prior-use in adults are the superior memory abilities of adults (Gathercole, 1999) or more flexible decision making in adults (Ernst, 2008). Children may have successfully acquired the prior but failed to integrate it with likelihood information (Dekker et al., 2015; Nardini et al., 2010). Our experiments do not allow us to distinguish between these possibilities.

For a theoretical account of our findings, we draw on the Reinforcement Learning literature. Model-based behavior leverages an understanding of the world’s structure to predict successful actions (Sutton and Barto, 1998). In order to use a model, the agent must have already learned the dynamics and causal structure of the environment, and our findings, along with others, suggest that the ability to do so is not fully developed in children (Decker et al., 2016). Increase in use of the prior during development and a decrease in simple biases may reflect a progression from use of a simpler model to a more complex one. Weaker cue combination in children can also be understood as a failure of model-based behavior, i.e. children may have undeveloped models of the behavior of real-world objects, and therefore lack understanding of when information should be integrated (Ernst, 2008; Gori et al., 2008; Nardini et al., 2010; Wei and Kording, 2012). Further experiments are needed to fully understand the change in learning strategies in sensorimotor integration over development.

There is great interest in understanding intelligent human behavior to build artificial systems with the same degree of functionality (Lake et al., 2016; Lecun et al., 2015; Marblestone et al., 2016). If our aim is to understand human behavior, we may benefit from understanding the process by which human abilities are learned in the first place. The algorithmic approaches of the future may benefit from implementing a development-like process, beginning with the kind of simple biases demonstrated in young children and progressing toward the flexible and complex model-based abilities of adults (Decker et al., 2016; Ullman et al., 2012).

If Bayesian computation is at the core of the neural code (Beck et al., 2008; Ma et al., 2006; Zemel et al., 1998), behavior should show the signatures of Bayesian inference under all conditions, including during development. Our results show that children do not use probabilistic information to the same extent as adults. The finding that children gradually shift toward the statistical optimum during development suggests that the brain learns to approximate Bayesian principles by means other than explicitly implementing Bayesian computations in neural circuits (Mandt et al., 2017; Rao, 2004). Our findings fit with ideas suggested by Jean Piaget on the role of constructivism in child development, i.e. that abilities are acquired through experience by building on more basic forms of knowledge (Piaget, 1954). In that sense, learning statistics may be seen as a very basic form of knowledge. While we may be born with a general learning architecture, it seems that statistics should not be seen as core knowledge (Spelke and Kinzler, 2007), but as an acquired skill.

